# A continuum of specialists and generalists in empirical communities

**DOI:** 10.1101/009985

**Authors:** Timothée Poisot, Sonia Kéfi, Serge Morand, Michal Stanko, Pablo A. Marquet, Michael E. Hochberg

## Abstract

Understanding the persistence of specialists and generalists within ecological communities is a topical research question, with far-reaching consequences for the maintenance of functional diversity. Although theoretical studies indicate that restricted conditions may be necessary to achieve co-occurrence of specialists and generalists, analyses of larger empirical (and species-rich) communities reveal the pervasiveness of coexistence. In this paper, we analyze 175 ecological bipartite networks of three interaction types (animal hosts-parasite, plant-herbivore and plant-pollinator), and measure the extent to which these communities are composed of species with different levels of specificity in their biotic interactions. We find a continuum from specialism to generalism. Furthermore, we demonstrate that diversity tends to be greatest in networks with intermediate connectance, and argue this is because of physical constraints in the filling of networks.

## Introduction

The functional diversity of ecological communities emerges through the simultaneous occurrence of species with contrasted resource use [1], habitat selection [2], and interactions [3,4]. Both empirical and theoretical studies have shown how different degrees of niche partitioning can promote functional diversity [5–7] and species persistence [8]. However, the co-occurrence of specialist and generalist species has received considerably less attention. The majority of studies seeking to understand the conditions for co-occurrence between populations of specialists and generalists in both biotic (*e.g.* predator–prey, host–parasite) and abiotic (*e.g.* habitat choice) interactions have focused on small communities [9–14].

Approaches based on model analysis or controlled experiments have two features impeding their generalization to large communities. First, the number of interacting organisms is often kept low, to facilitate model analysis or because of experimental constraints. Studies investigating the co-occurrence of species with contrasted specificities assume no intermediate situations between the endpoints of specialism and generalism, whereas natural systems exhibit a continuum [15,16]. Second, it is unclear to what extent results can be scaled up to more realistic communities. Stouffer and colleagues [17] showed that because adding species and interactions increases the potential for complex population dynamical feedbacks, complete, realistic networks tend to exhibit different behaviors than simple modules (*i.e.* those typically used in models or experiments), begging for an analysis of co-occurrence in empirical communities.

Network theory offers powerful tools to describe ecological communities [18] and the distribution of species specificity within them [19]. In a species interaction network, each species is a node, and each interaction is an edge, connecting a pair of nodes. From a network perspective, one can measure specificity by counting the number of links it has with other species (its degree), or by measuring aspects of the distribution of the strengths of such links [19]. Previous work described the degree distribution (*i.e.* the distribution of how many interactions each species establishes and receives) of empirical networks, and revealed a continuum from highly specialized to generalists species [20]. While much is known about the factors (*e.g.* biotic [21], abiotic [14,15], developmental and physiological [22]) driving the specialization of single species, less is known about the spectrum of specificities and niche-overlaps that can co-occur in large ecological networks, and reasons for different spectra. As the co-occurrence and interactions between specialized and generalized species is key to maintaining functional diversity [23], promoting community stability [24], and ensuring network persistence [3], there is a need to investigate the extent and properties of this co-occurrence.

In a previous paper [1], we argued that the specialisation of different types of interactions is likely to be shaped by the same set of core mechanisms, expressed in a different ways or with different intensities. At the community level, this leads to the expectation that the same relationships between specificity, the co-occurrence of specialists and generalists, and other metrics of community structure would occure for different types of ecological interactions, despite different types of networks, dominated by positive or negative interactions, occupying different parts of this gradient [25]. In this study, we use a dataset of interaction networks spanning three contrasted types of ecological interactions (herbivory, parasitism, and mutualism), to characterize the extent to which species with different specificities can co-occur within the same community. In line with our expectation and past empirical data, we find a continuum from networks of mostly-specialized to mostly-generalized species, with the potential for specialist/generalist co-occurrence being greater at intermediate connectance. One central result is that empirical data show consistently more variation in specificities of all species on the upper network level (parasites, herbivores, pollinators; hereafter called “strategy diversity”) than predicted by two contrasting null models. This suggests (i) that organisms with very different levels of specificity can co-occur in most natural systems, and (ii) that ecological or evolutionary mechanisms are acting to maintain high diversity in the range of specificities.

## Methods

### Datasets

We employ three datasets: two for antagonistic (ectoparasite–animal host and insect herbivore–plant) interactions, and one for mutualistic (pollinator–plant) interactions. Parasitism networks were from Stanko and colleagues [26,27] and consist of 121 networks of ectoparasites infecting rodents in Central Europe, collected in a range of continental ecosystems over a period of 19 years. Herbivory networks (a total of 23) were collected by Thébault and Fontaine [25] from various literature records. Data on mutualistic interactions are the 29 “plant–pollinators” networks deposited in the *InteractionWeb* database (http://www.nceas.ucsb.edu/interactionweb/) as of May 2012. These data are insect–plant contacts, aggregated from different sources, spanning a period of over 30 years. Species with no interaction were removed from the original datasets. Some networks had less than 1000 possible randomizations, which did not allow for efficient or meaningful randomisation [28], and as such were discarded from the analysis. The final dataset has 115 parasitism networks, 6 herbivory networks, and 12 pollination networks. Because the sample size is unbalanced, we put particular emphasis on the discussion of parasitism networks.

### Network analyses

Each bipartite network is represented by its adjacency matrix **M** with *T* rows (for the upper level, *i.e.* ectoparasites, herbivores, and pollinators) and *L* columns (for the lower level, *i.e.* animal hosts and plants being consumed or pollinated). In each network, **M**_*ij*_ represents the existence of an interaction between species *i* and species *j* [29]. For each network, we calculate its size (*Z* = *L* × *T*), and connectance (Co, the proportion of established interactions). We focus our analyses on the upper level, since we have more knowledge of specialization mechanisms for these organisms [30]. Nestedness, a measure that reflects whether specialist species interact with the same species as generalists, is calculated using the NODF (Nestedness based on Overlap and Decreasing Fill) measure [31]. NODF is insensitive to network asymetry (the relative number of species at each of the two levels) and size. Modularity measures the extent to which species form well defined, densely connected, groups, with few connections between groups. Modularity is estimated using the LP-BRIM method [32], which both increases detection compared to the adaptive BRIM method, and is less computationally intensive [33]. For each network, we retained the highest modularity *Q*_*bip*_ [34] observed in a total of 1000 replicate runs.

We contrast empirical observations with the predictions of two different null models, each based on the impact of different aspects of network structure. For each null model, we filled a network through a Bernoulli process, in which the probability of each pairwise species interaction occurring (P_*i* *j*_) is determined in one of the following ways. Null model I [35] is connectance based and assigns the same probability to each interaction, P_*i* *j*_ = *Co*. Compared to the empirical network on which they are based, simulated networks can have the same connectance, but a potentially different degree distribution. Null model II [3] uses information about species degree (the number of interactions established/received) to calculate the probability that a particular interaction will occur. This probability is P_*i* *j*_ = (*T* × *G*_*i*_ + *L* ×*V*_*j*_)/(2 × *Z*), where *G*_*i*_ and *V*_*j*_ are, respectively, the generality (number of interactions) of upper level species *i*, and the vulnerability (number of interactions) of lower level species *j* [36]. Simply put, the probability of the interaction occurring is the mean of the degrees (ranged in 0–1) of the two species involved. Note that the first null model is nested into the second.

Each of these models was applied to each network in the dataset, so as to generate 1000 random networks (meaning that each empirical network was fed into the model to generate a total of 2000 randomizations). Each of these networks was analyzed using the same methods as for empirical networks.

### Quantifying specificity

We quantify specificity based on the proportion of available species with which a focal species interacts [37], using a ranged version of Schoener’s generality. For each species *i* of the upper level (*e.g.* parasites), its specificity is given by

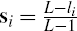

where *L* is the number of lower level species (*e.g.* hosts) found in the network, and *l*_*i*_ is the number of interaction partners of species *i*. The vector s is the distribution of specificities at the network scale.

Values of 1 indicate complete specialism (single partner), and values of 0 indicate complete generalism (all possible partners).

### Quantifying strategy diversity

We quantify two aspects of the co-occurrence of specialists and generalists (*i.e.* “strategy diversity”). First, “specificity range” or *R*, is simply the difference between the specificity of the most and least specialized organisms, such that

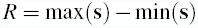

*R* is maximized when at least one completely specialized species *k* (s_*k*_ = 1) is found in the same network as one (or more) completely generalized species *l* (s_*l*_ = 0).

A second measure of the distribution of specificities within a network is its evenness, denoted *E*. We define s′ as all the unique values of s, rounded to the second decimal place. We define *U* as the ordered set of s′ values and *u* as each of the elements of this set. Thus *p*(*u*) is the probability associated to a given element of *U*. For example, if s′ = [0.1, 1, 1, 0, 0.4], then *U* = [0, 0.1, 0.4, 1], *p*(*u* = 1) = 2/5, and *p*(*u* = 0) = 1/5. With this information, we calculate the self-information [38] of *u* as *I*(*u*) = -ln(*u*), and based on these two sets of values, we calculate the Shannon’s entropy of the distribution of specificity values as

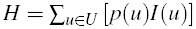

If *U* takes on *N* possible values, then the theoretical maximum of *H* (attained when all values of s′ are unique, *i.e.* no two species are equally specialised) is

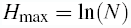

To eliminate any scaling effect that might occur due to different network sizes, we take the exponentials of these values [39], such that the standardized value of *E* is

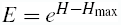

It follows that *E* = 1 when no two organisms have the same level of specificity, and *E* = 0 when all values of s′ are equal. Note that rounding to the second decimal place allows accounting for the fact that some organisms may have very similar (but not exactly equal) specificities. Small differences in the values of specificity are less important than the potential amplitude of measurement error, as preliminary tests indicated that the rounding of s′ does not qualitatively change observed relationships. It is also known that small differences in link strength have little or no impact in larger networks [40].

Finally, we present a simple summary statistic that we call “strategy diversity” (*D*),

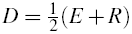

which given that both *E* and *R* take values in [0, 1], will also return values in this range. *D* = 1 indicates that the specificity values found in a network range from highly specialized to highly generalized *and* are evenly distributed. *D* = 0 means that a network is composed entirely of species sharing the same specificity values. The two advantages of *D* are (i) it accounts both for the range of specificities and their distribution, and (ii) it is independent of the observed specificity values. We expect strategy diversity (*D*) to peak at intermediate values of connectance and specificity, to increase with nestedness, and to decrease with modularity (Fig. 1). The reasoning is as follows. Interaction matrices are physically constrained objects, in that adding interactions will modify their properties, and thus produce artifacts [28,41]. By definition, a perfectly nested network maximizes strategy diversity [31], and a modular network tends to minimize it. A matrix with minimal fill for a given size has all interactions on the diagonal, and is therefore highly specialized, with no strategy diversity. Conversely, a completely filled network is extremely generalized, and thus has no strategy diversity.

**Figure 1:**
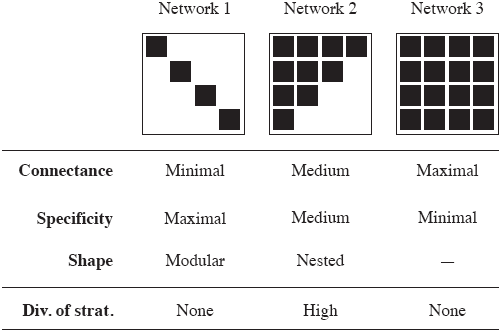
Expected relationships between connectance and other metrics.

## Results

All types of network tend to have more strategy diversity and to be composed of more specialized species than expected by chance (Table 1). For each empirical network, we measure whether its structural properties (strategy diversity, nestedness, modularity) are higher or lower than expected by chance using the two null models. Our results are reported in Table 1. Both null models gave consistent results regarding whether the empirical networks represented a deviation from random expectations. Host-parasite networks are on average less modular than expected, herbivory networks are more, and there is no clear trend in pollination networks. There is a marked tendency towards higher than expected nestedness in all types of interactions.

**Table 1.** Results of the null models analyses. For each network metric, and for each null model, we indicate the proportion of networks that had significantly larger or smaller values than expected by chance. A network has a significantly different value from the prediction when the empirical value falls outside of the 95% confidence interval for the value as mesured on randomized networks [55]. NS: no significant difference in stategy diversity. *D*: strategy diversity. *S*: average specificity.

Figure 2 presents the distributions of specificity, connectance, nestedness, and modularity in networks that are either more or less functionally diverse than expected under the assumptions of null model II (using the outcomes of model I yields the same qualitative results; see Table 1). Regardless of the baseline differences between types of network for each of the metrics considered, higher diversity responded in a consistent way to variation in the other metrics. Networks with higher average specificity tended to have lower average strategy diversity, higher connectance, higher nestedness, and lower modularity (Table 2). There are significant interactions between all of the variables and the network having higher strategy diversity than expected by chance, with the exception of modularity (Table 3). These four metrics alone account for 96% of the variance of strategy diversity, and 63% of the variance in the deviation of this same metric. All metrics except modularity had a significant impact on strategy diversity. Interestingly, connectance was the best predictor of strategy diversity, whereas nestedness was the best predictor of the extent to which strategy diversity in the empirical networks deviates from random expectations. This is because by definition, null model testing removes most of the effects of connectance. The type of ecological interaction was not significant; detecting possible significance would have probably required a larger sample size for non-parasitic networks.

**Figure 2:**
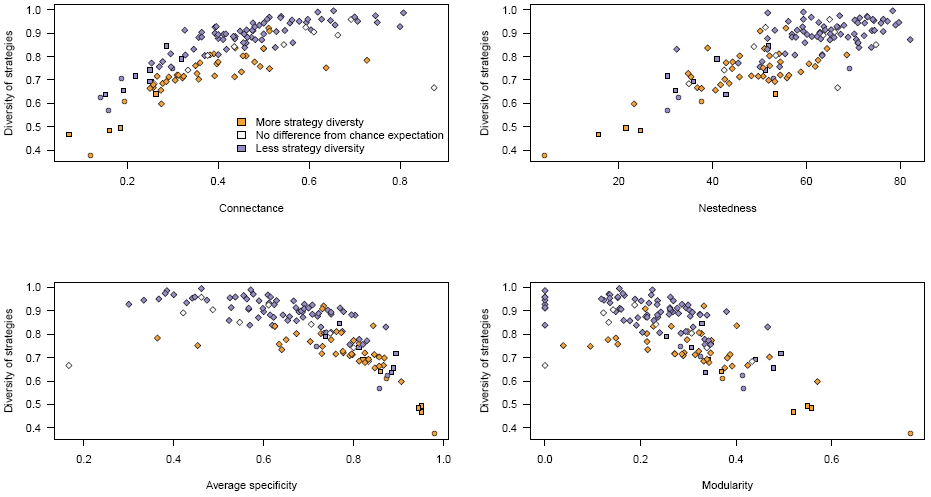
Values of average specificity, nestedness, connectance, and modularity for networks with more (orange) or less (purple) strategy diversity than expected by chance. The results within a type of interaction are all highly consistent. For this analysis *only*, networks that were as functionally diverse as expected (as determined by the Null Models) were removed, since their strategy diversity can be explained solely by either their connectance or degree distribution. Types of interaction are given on the x axis, with networks separated as a function of whether they have more (orange) or less (purple) strategy diversity than expected by chance (under the assumptions of the second, more restrictive null model).

**Table 2.** Analysis of the results presented in Fig. 2. We used a two-sample t-test to determine differences from chance expectations for networks with either less, equal, or more strategy diversity. We observe that all metrics are different from chance expectatons for parasitism networks, but not for other interaction types (although our failure to report an effect is most likely due to the small sample size, as indicated by certain large confidence intervals).

**Table 3.** Analysis of variance partitioning (ANOVA on linear additive models) of the effects of connectance, nestedness, mean specificity, and modularity, on strategy diversity, and the excess strategy diversity (deviation of empirical values from simulated networks as asssessed by the Null Model analysis). Preliminary analyses showed no impact of the interaction type on these relationships, so this factor was not included as a covariate. *D*: strategy diversity.

We finally examine the relationships between network metrics and strategy diversity (Fig. 3). Strategy diversity increases with connectance (it is expected to be 0 for a connectance of 1, but no network in our dataset is densely connected), decreases with average specificity (as before, strategy diversity is 0 if mean specificity is 0), increases linearly with nestedness, and decreases with modularity. An interesting result in this analysis is that the trend is the same for all three types of interaction considered, with the exception that herbivory and pollination networks tended to occupy the “low connectance” end of the gradient; they behave in the same way as do parasitism networks, reinforcing the idea that structural constraints such as that introduced by connectance may be driving emergent network properties [28,42].

**Figure 3:**
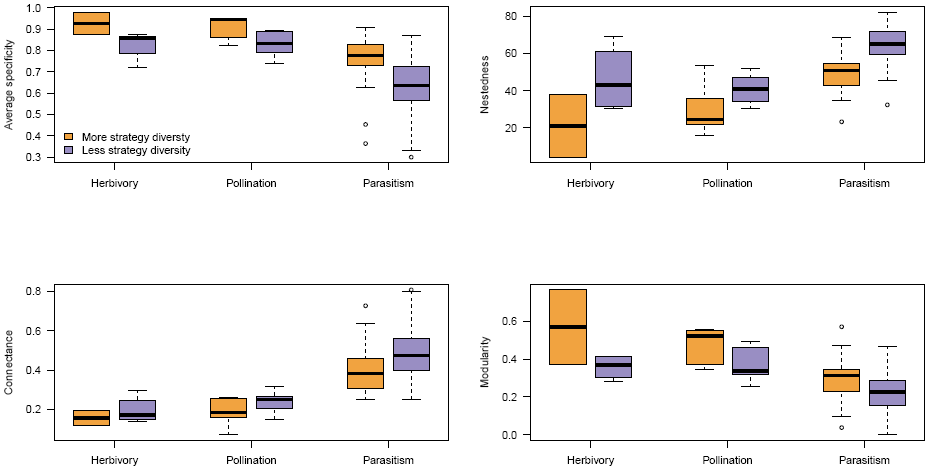
Scatterplot of strategy diversity versus other network metrics. Regardless of the interaction type, strategy diversity responds in a similar way to other network metrics. Points are colored as in Figure 2. Triangles are host-parasite systems, squares are plants-herbivores, and circles are plants-pollinators. Empty triangles are host-parasite networks that have as many strategy diversity as expected.

## Discussion

Several mechanisms have been proposed to explain the co-occurrence of potentially competing species, including behavior [43], spatial or temporal heterogeneity [9], and trade-offs associated with species interactions [1,12,44]. Ecological factors such as environmental and spatial heterogeneity and evolutionary processes such as niche partitioning may permit the coexistence between competing species with similar and/or different number of resources [45,46]. However, most of these results were obtained in systems of low complexity, and the extent to which specialists and generalists co-occur in natural communities remains to be evaluated. By analyzing three bipartite network datasets covering a range of both ecological and structural situations, we show how co-occurrence can be linked with other topological network properties. This calls for a better integration of network methodology to the analysis of community structure, with the aim of understanding the co-occurrence of species with different specificities.

Most emergent network properties could be predicted based on connectance alone [28]. This included, notably, components of the degree distribution (how many interactions are established/received by each species) involved in determining nestedness. The fact that the relationship between connectance, emergent metrics (such as nestedness and modularity), and strategy diversity is conserved across types of ecological interactions can be explained in part by these physical constraints. The fact that some interactions appear more or less specialised reflects average differences in connectance in these communities. Null models analysis nonetheless reveals that, for all types of interaction, approximately two-thirds of all networks had *more* strategy diversity than expected by chance; this suggests that despite physical constraints, ecological and/or evolutionary mechanisms are involved in promoting high diversity [8,47].

Overall, we report that networks with higher nestedness and lower modularity, also had more strategy diversity than expected under the assumptions of the two null models. If the main difference between interaction types is their connectance, then the different mechanisms involved must be studied alongside their impacts on network structure. Species specialization is regulated by differences in life-history traits [1], competition for access to resources [45,48], or phylogenetic conservatism in attack/defense strategies [49]. Through their impact on species range of resources used, these factors are likely to be involved in driving network structure, and connectance in particular. For example, in herbivorous systems, plants may employ multiple defenses against enemies, including the release of toxic compounds [50] and/or attraction of a herbivore’s natural enemies [51–54]. The simultaneous existence of different levels of defense such as those mentioned above may promote lower connectance. It can also result in the faster diversification of exploitation strategies at the upper level (in the sense that enemies specialize on a *defense mechanism* rather than on the set of defended species) than in other types of interaction rely on narrower sets of mechanisms [15]. This may result in the maintenance of high strategy diversity relative to connectance in some antagonistic interactions.

In summary, although the ecological nature of an interaction (mutualistic or antagonistic) has an impact on network structure, higher than expected strategy diversity appears to be a conserved property in bipartite ecological networks. The particular position occupied by a network along a continuum of, *e.g.* connectance or nestedness, can emerge because of the life-history traits of species establishing interactions, and we suggest that increased attention should be given to understanding how fine-scale mechanisms at the individual or population level drive the structure of community-level networks. It is nonetheless clear that despite theoretical predictions, generalists and specialists are often found together in nature. Understanding this gap between predictions and observations will be a major challenge that should be addressed by investigating the mechanisms of coexistence and co-occurrence in large multi-species communities.

## Acknowledgments

We thank E. Canard, V. Devictor, I. Gounand, S. Fellous & N. Mouquet for comments, and the Canadian Research Chair on Continental Ecosystems Ecology for providing computational support. We thank É. Thébault and C. Fontaine for sharing data for plants and herbivores systems. The bipy package used for the analyses, and the C99 program used for the generation of random networks under the three null models are available at http://github.com/tpoisot/bipy and http://github.com/tpoisot/CNullModels. MS was funded by Slovak Research and Development Agency grant No. 0267-10. TP was support by a CRD grant from NSERC, and a PBEEE post-doctoral scholarship from FQRNT/MELS. MEH is funded by grants from Agence Nationale de la Recherche ‘EvolStress’ (ANR-09-BLAN-099-01), the CNRS (PICS06313) and the McDonnell Foundation (JSMF 220020294/SCS-Research Award).

